# Neglectful maternal caregiving involves altered brain plasticity in empathy-related areas

**DOI:** 10.1101/654434

**Authors:** María José Rodrigo, Inmaculada León, Lorna García, Juan Andrés Hernández-Cabrera, Ileana Quiñones

## Abstract

The maternal brain undergoes functional and structural adaptations to sensitive caregiving that are critical for ensuring infant wellbeing. This study investigates brain structural alterations in neglectful caregiving and their impact on mother-child interactive behavior. High-resolution 3D volumetric images were obtained on 25 neglectful (NM) and 23 non-neglectful control (CM) mothers. Using Voxel-Based Morphometry, we compared gray and white matter volume (GMV/WMV) between the two groups. Mothers also completed an empathy scale and participated with their children in a standardized play task (Emotional Availability Scale, EA). NM, as compared to CM, showed GMV reductions in right insula, anterior/middle cingulate, and right inferior frontal gyrus (IFG), as well as WMV reductions in bilateral frontal regions. A GMV increase was observed in the right fusiform and cerebellum. Regression analyses showed a negative effect of fusiform GMV and a positive effect of right frontal WMV on EA Mediation analyses showed the mediating role of emotional empathy in the positive effect of insula and IFG, and the negative effect of cerebellum on EA. Neglectful mothering involves an altered plasticity in emotional empathy-related areas and in frontal areas associated with poor mother-child interactive bonding, indicating how critical the structural changes in these areas are for infant wellbeing.

## Introduction

The experience of being a mother involves functional and structural brain changes supporting the establishment of sensitive caregiver responses to infants (Kim, Strathearn, & Swain, 2016). Mothers compared to non-mothers exhibited a specific right prefrontal response when discriminating infant facial expressions (Nishitani, Doi, Koyama, & Shinohara, 2011). Mothers also respond to their own versus unfamiliar infants’ cry sounds and faces, with a more intense activation in regions involved in viso-emotional processing, empathy or emotion regulation (Kim et al., 2016; Rocchetti et al., 2014). A higher activation is also found when mothers are responding to an emotional face-matching task in the late as compared to the early postpartum period, which may reflect long-lasting adjustments in emotional empathy-related areas (Gingnell et al., 2015). Moreover, some of these areas also undergo structural increases in gray matter during the postpartum period, suggesting maternal brain adaptations to the child’s evolving needs (Barba-Müller, Craddock, Carmona, & Hoekzema, 2018). From this perspective, it is reasonable to expect alterations in the brain adaptations in those mothers exhibiting a severe lack of response to their own child’s needs. The current study addressed this new topic by investigating possible structural alterations in the maternal brain underlying a drastic disregard of the infant’s basic needs, and their functional association with mother-child interactive behavior. Finding this association may further illustrate how critically adaptive these structural changes are for infant wellbeing.

An example of extremely insensitive caregiving is maternal neglect that consists of the mothers’ failure to provide the child for food, clothing, shelter, medical care, supervision or emotional support. It is the most common and severe form of child maltreatment that puts the child’s safety at risk (Petersen, Joseph, & Feit, 2013; Stoltenborgh, Bakermans-Kranenburg, & van IJzendoorn, 2013). Negligence also disrupts the establishment of a child’s secure attachment and healthy psychosocial development (Weinfield, Sroufe, Egeland, & Carlson, 2008), and entails negative and cumulative behavioral and neurobiological alterations for the offspring (see review by Teicher, Samson, Anderson, & Ohashi, 2016). Studying the neurobiological basis of maternal neglect could help in the tuning of intervention efforts to reduce risk and promote infant health.

Our first aim was to examine gray and white matter volume (GMV/WMV) differences between neglectful (NM) and non-neglectful control (CM) mothers who are sociodemographically similar. Searching for brain adaptations that presumably fail to occur in NM as compared to CM, we expected that at least part of the volumetric differences would overlap with empathy-related areas. Those areas undergo functional and structural adaptations to sensitive caregiving in normal mothering (Kim et al., 2016). Parental empathy, defined as the appropriate perception, understanding, and experience of an infant’s emotional states, is one of the crucial abilities for providing caring responses to one’s own infant’s needs, that is lower in NM (León et al., 2014; Rodrigo et al., 2011). NM’s lower empathic skills seem to be tightly coupled with their lower brain reactivity to infant cues. NM showed a generic attenuated response to infant and adult crying faces in viso-limbic areas, such as bilateral lingual, bilateral cerebellum, bilateral fusiform gyrus, right hippocampus, parahippocampal gyrus, and right amygdala. Reduced activations are also specifically shown to infant crying faces in left middle frontal and anterior cingulate areas, underscoring their difficulties to respond to the infant’s emotional cues (León et al., in press). Brain differences were assessed using Voxel-Based Morphometry (VBM; Ashburner & Friston, 2000), an approach used to quantify structural brain properties for the investigation of volume differences in brain anatomy.

With respect to the GMV structural changes, adults with higher scores in the emotional Empathic Concern scale, defined as the embodied simulation of and sympathy with others’ emotions, showed increased GMV in the anterior insula (Eres, Decety, Louis, & Molenberghs, 2015; Mutschler, Reinbold, Wankerl, Seifritz, & Ball, 2013) and the inferior frontal gyrus (Banissy, Kanai, Walsh, & Rees, 2012) than those with lower scores. In turn, adults with higher scores in the cognitive Perspective Taking scale, defined as the mental-state understanding and perspective taking from others, showed increased GMV in the middle cingulate cortex and the adjacent dorsomedial prefrontal cortex than those with lower scores (Eres et al., 2015). Importantly, similar increases in maternal brain volume were also reported as an adaptation to parenting. GMV increases in sensitive mothers with perceived higher parental care in childhood was also found in the superior/middle frontal areas, orbitofrontal gyrus, superior temporal gyrus, and fusiform gyrus (Kim et al., 2010). Given all these findings, we expected to find some volumetric differences in NM compared to CM in empathy-related areas, given their severe disregard of the child’s needs and their lower scores in empathic concern (León et al., 2014; Rodrigo et al., 2011).

Structural alteration in WMV is also expected in NM. WM disruption has been reported in postpartum depression, in the fronto-thalamic circuit and in interhemispheric connectivity (Silver et al., 2018). In turn, a higher Empathic Concern in adults has shown to be positively associated with greater WM structural integrity in the tracts linking areas involved in affective processing of infant faces, such as the inferior longitudinal fasciculus (ILF) and the inferior fronto-occipital fasciculus (IFOF), (Parkinson & Wheatley, 2012). Given that neglectful mothers showed a volume reduction in the ILF and IFOF in a Diffusor Tensor Imaging study (Rodrigo et al., 2016), we also expected a decrease in WMV in those areas traversed by these tracts in relation to CM.

As a second aim, we examined the functional impact of GMV and WMV alterations on Emotional Availability (EA), as a proxy for the quality of mother-child bonding interactions. EA measures through a play task the ability to read and respond appropriately to each other’s emotional signals in mother-child dyads (Biringen, 2000; Biringen, Derscheid, Vliegen, Closson, & Easterbrooks, 2014). EA is also predictive of the mothers’ reported child attachment (Altenhofen, Clyman, Little, Baker, & Biringen, 2013). A previous study had shown lower scores in NM than in CM in EA associated with a volume reduction in the ILF and IFOF (Rodrigo et al., 2016). Therefore, we expected to find a positive association between GMV and WMV alterations and lower EA scores in NM.

Once the possible impact of GMV and WMV alterations on maternal sensitivity in NM had been tested, we wanted to go a step further determining the possible role of trait emotional and cognitive empathy in that relation. A higher score in Emotional Concern has been associated with the intention to provide care to the needed child (Lin & McFatter, 2012) and the expression of maternal warmth and positive affect (Stern, Borelli, & Smiley, 2015). In turn, a higher score in Perspective Taking has been related to increases in one’s own infant’s pupil dilation in response to others’ emotional displays, as an early precursor of empathy (Upshaw, Kaiser, & Sommerville, 2015). Therefore, if volumetric group differences are found in areas corresponding either to the emotional or cognitive empathy circuits, we expected that introducing Empathic Concern or Perspective Taking, respectively, as mediators would increase the possibilities of finding an association between volumetric differences and EA.

Altogether, this study can provide evidence of the morphometric alterations underlying neglectful mothering and the role played by the emotional and cognitive empathy-related areas in maternal caregiving.

## Methods

### Participants

Forty-eight mothers (25 NM and 23 CM) voluntarily participated in the experiment. They all were recruited through the same Primary Health Center in Tenerife, Spain. Written consent was obtained from all the participants. The Ethics Committee of the University of La Laguna approved the study’s protocol. General inclusion criteria were being the biological mother of a child under three years old who had not been placed in foster care at any point in their history nor been born prematurely or suffered perinatal or postnatal medical complications, according to the pediatricians’ reports. Specific inclusion criteria for the neglectful mother group were a substantiated case of neglect registered in the last 12 months by Child Protective Services (CPS), and complying with the indicators of the Maltreatment Classification System (MCS) for severe neglect (Barnett, Manly, & Cicchetti, 1993). Thus, these mothers scored positively on physical neglect (inadequate food, hygiene, clothing, and medical care), lack of supervision (child is left alone or in the care of an unreliable caregiver), and educational neglect (lack of cognitive and socioemotional stimulation and lack of attention to the child’s education). Inclusion criteria for the control group were negative scores in all the MCS neglect indicators, and the absence of CPS or Preventive Services records for the family. As for the sociodemographic profile of mothers, they were all in their early 30s, had a similar number of children and mean age of the target child, and shared similar, low socioeconomic backgrounds (Table 1). According to the neglect risk profile rated by the social workers on presence /absence of risk indicators for neglect, most mothers in the neglectful group had a history of childhood maltreatment or neglect (of the mother when she was a child). They also scored positively in poor household management, disregard of the child’s needs and rigid/inconsistent discipline norms [see Appendix 1 for details on the risk profile measures].

**Table 1.** Sociodemographic and neglect risk profile in Neglectful and Control Groups

| | Neglectful group<br>(*n* = 25)<br>*M* (*SD*) or % | Control group<br>(*n* = 23)<br>*M* (*SD*) or % | *F*(1,42)/<br>$\chi^2$ |
| --- | --- | --- | --- |
| Sociodemographic profile |  |  |  |
| Mean age of mother | 29.2 (7.0) | 33.43 (3.4) | -2.71** |
| Number of children | 2.08 (0.8) | 1.65 (0.6) | 1.93 |
| Mean age of the target child | 2.8 (1.5) | 2.1 (1.8) | 1.5 |
| Rural areas (%) | 45.8 | 37.5 | 0.61 |
| Level of education (%): |  |  | 2.93 |
| Primary | 72 | 47.0 |  |
| Secondary school | 16 | 30.0 |  |
| > Secondary school | 12 | 21.7 |  |
| Employment (%) | 44 | 34 | 5.32 |
| Neglect risk profile |  |  |  |
| History maltreatment/neglect (%) | 68 | 17.4 | 10.49*** |
| Intimate partner conflict (%) | 5 | 0 | 3.68 |
| Chronic physical illness (%) | 4 | 0 | 2.51 |
| Poor household management (%) | 84.2 | 0 | 24.29*** |
| Disregard health/education needs (%) | 57 | 0 | 12.79*** |
| Disregard emotion/cognitive needs (%) | 89 | 0 | 27.24*** |
| Rigid/inconsistent norms (%) | 68 | 0 | 16.83*** |

\*\**p*<.01; \*\*\**p*<.001.

### Behavioral and personality measures

#### The Mini International Neuropsychiatric Interview

(M.I.N.I. 6.0; Spanish version (Ferrando, Bobes, Gibert, Soto, & Soto, 2000), including 15 major psychiatric disorders, was applied (Table 2). The two groups mainly differed in five psychopathological variables marked in italics that survived the Bonferroni correction, which were submitted to a Principal Component Analysis. Results gave a one factor solution: “Psychiatric Disorders”, with moderate inter-correlations among the five variables, KMO = 0.68, Eigenvalue = 2.74, with an explained variance of 55%, with the coefficient scores in Psychiatric Disorders being higher in neglectful (*M* = 0.63, *SD* = 1.02) than in control mothers (*M* = −0.69, *SD* = 0.21), *t*(46) = 6.3; *p* = .000; *δ* = 1.82. None of the mothers in either group was being medicated for psychiatric disorders at the time of testing. The coefficient score in Psychiatric Disorders (PD), was used as a regressor in the SPM model to control as much as possible for its effect on brain volume differences. Given that the dichotomized Group of mothers and PD were certainly related (r = 0.67), we previously rule out their potential collinearity before its inclusion in the model as a covariate [See Appendix 2, Table A1]. All the analyses were performed with R (R Core Team, 2017).

**Table 2.** Psychopathological conditions stratified by Group

| | Neglectful group<br>(*n* = 25)<br>*M* (*SD*) | Control group<br>(*n* = 23)<br>*M* (*SD*) | *t*(46) | Effect<br>size<br>$\delta$ |
| --- | --- | --- | --- | --- |
| *Major Depressive Episode* | 2.0 (2.6) | 0.2 (0.5) | 3.36** | 0.97 |
| Dysthymia | 1.6 (2.2) | 0.3 (0.5) | 2.75** | 0.79 |
| Suicidality | 0.5 (0.8) | 0 | 2.68* | 0.81 |
| *Hypo/Manic Episode* | 1.8 (2.1) | 0.1 (0.3) | 3.94** | 1.14 |
| *General Panic Disorder* | 6.8 (5.7) | 0.7 (2.1) | 4.93*** | 1.42 |
| Agoraphobia | 0.7 (1) | 0.2 (0.4) | 2.33* | 0.70 |
| Social Phobia | 0.6 (1) | 0 | 2.50* | 0.75 |
| Obsessive-Compulsive | 1.2 (1.6) | 0.2 (0.6) | 2.46* | 0.74 |
| Post-traumatic Stress<br>Disorder | 1.6 (2.6) | 0.9 (1.8) | 0.96 | 0.29 |
| Alcohol<br>Dependence/Abuse | 0.1 (0.2) | 0.2 (0.5) | 0.78 | 0.23 |
| Drug Dependence/Abuse | 0.2 (0.4) | 0 | 1.89 | 0.57 |
| Psychotic Disorders | 0.7 (1.4) | 0.2 (0.5) | 1.68 | 0.49 |
| Bulimia Nervosa | 0.1 (0.2) | 0 | 1 | 0.29 |
| *Generalized Anxiety*<br>*Disorder* | 3.4 (3.4) | 0.6 (0.9) | 3.76*** | 1.08 |
| *Antisocial Personality* | 1.2 (1.2) | 0.1 (0.3) | 4.08*** | 1.18 |

\* $p \leq 0.05$ ; \*\* $p \leq 0.01$ ; \*\*\* $p \leq 0.001$ ; Variables with italic font were submitted to the Principal Component Analysis.

#### Emotional Availability

*(EA)* was measured in the context of mother-child free play using the EA Scale: Infancy to Early Childhood Version (Easterbrooks & Biringen, 2005). This scale operationalizes parental and child behavior in six scales that were factorized into one factor (EA) using Principal Component Analysis, given the existence of high inter-correlations among the scales. Two external observers blind to the mothers’ grouping made the ratings from the videos, and the inter-rater reliability of the ratings in each scale was adequate [see Appendix 3 and Table A2 for the scales and testing]. EA was lower in neglectful (*M* = −0.61, *SD* = 0.92) than in control dyads (*M* = 0.66, *SD* = 0.55), *t*(46) = −5.75; *p* = .000; *δ* = 1.66 and it was used as a dependent variable.

#### Interpersonal Reactivity Index

(IRI; Spanish version (Pérez-Albéniz, De Paúl, Etxeberría, Montes, & Torres, 2003). Only the Empathic concern (EC) and the Perspective Taking (PT) scales were used. EC taps the respondents’ feelings of warmth, compassion, and concern for others (emotional empathy), whereas PT describes the tendency to spontaneously adopt others’ points of view in everyday life (cognitive empathy). The scores in EC and PT were lower in NM (*M* = 26, *SD* = 3.59; *M* = 23.56, *SD* = 4.35) than in CM (*M* = 28.08, *SD* = 3.56; *M* = 26.21, *SD* = 3.74), *t*(46) = −2.01; *p* = .05; *δ* = 0.58; *t*(46) = −2.25; *p* = .05; *δ* = 0.65.

### MRI Processing and Analysis

#### Structural Image Acquisition

High-resolution T1-weighted MPRAGE anatomical volumes were acquired on a General Electric 3T scanner. A total of 196 contiguous 1mm sagittal slices were acquired with the following parameters: repetition time = 8.716 ms, echo time = 1.736 ms, field of view = 256 × 256 mm^2^, in-plane resolution = 1 mm × 1 mm, flip angle = 12.

#### Voxel-Based Morphometry Processing

T1 images were pre-processed using the Voxel-Based Morphometry (VBM) toolbox (http://dbm.neuro.uni-jena.de/vbm.html) and the SPM8 software package. Images were corrected for bias-field inhomogeneity; tissue-classified into gray and white matter and cerebrospinal fluid; and registered to standard space using high-dimensional DARTEL normalization (Ashburner, 2007). The segmentation approach used is based on an adaptive Maximum A Posterior technique, which does not need a priori information about tissue probabilities (Rajapakse, Giedd, & Rapoport, 1997). This procedure was further refined by accounting for partial volume effects and by applying a hidden Markov random field model which incorporates spatial prior information of the adjacent voxels into the segmentation estimation (Tohka, Zijdenbos, & Evans, 2004). To measure regional differences in absolute GMV and WMV in the obtained volumetric segmentations, the warped images were modulated. All the normalized-modulated images were smoothed with a filter of a 10 mm Gaussian kernel. An additional quality check based on the sample homogeneity was conducted.

#### Statistical Analysis of GMV and WMV

GLM analyses (i.e., in SPM, Full Factorial Design, (Gläscher & Gitelman, 2008) were performed using the individual gray/white matter volumetric segmentations as dependent variables and including Group (Control vs. Neglectful mothers) as a between-subject factor. The models included one regressor described above as *Psychiatric Disorders.* The age of the mothers (mean centered) was also included as a nuisance covariate.

The resulting statistical parametric maps were thresholded at a peak level of *p* < 0.001 (uncorrected) and then the cluster spatial extent adjusted to capture only those clusters corrected for multiple comparisons using the whole-brain or the small-volume FWE corrections (*p* < 0.05). All local maxima were reported as MNI coordinates. Gray matter regions were labeled with reference to the Automated Anatomical Labeling atlas (AAL; Tzourio-Mazoyer et al., 2002)

## Results

### Altered GMV and WMV in neglectful mothers

Voxel-based morphometry analysis showed three distributed clusters with a pattern of reduced GMV in NM than in CM. Each of these clusters spanned several contiguous anatomical regions in both hemispheres (Table 3 and Figure 1). Cluster 1 included a large midline region comprising the anterior/middle cingulate (ACC/MCC) cortex with extension to both hemispheres, superiorly to the left precentral gyrus, the left superior frontal gyrus and the most inferior part of the right supplementary motor area. Cluster 2 involved a broad region from the pars triangularis within the inferior frontal gyrus to the adjacent part of the middle frontal gyrus. Cluster 3 was circumscribed to the posterior part of the right insula. In addition, VBM analysis showed one posterior cluster exhibiting the opposite pattern, that is, increased GMV in NM. As this cluster comprised a broad region it was anatomically divided into two distinct areas, fusiform and cerebellum, to test their effects separately.

**Table 3.** Gray and white matter volumetric differences showing cingulate, frontal and insula reduced GMV clusters (NM < CM), one posterior increased GMV cluster (NM > CM) and one frontal reduced WMV cluster (NM < CM).

|  | Coordinates<br>{mm} |  |  | Cluster-level |  | Peak level |
| --- | --- | --- | --- | --- | --- | --- |
|  | x | y | z | p values<br>(FWE-corr) | Cluster size<br>(voxels) | T scores |
| **Neglectful < Control Mothers (GMV)** |  |  |  |  |  |  |
| Cingulum_Mid_R | 6 | 38 | 32 | 0.000 | 7776 | 5.804 |
| Cingulum_Mid_L | -6 | 15 | 36 |  |  | 5.227 |
| Cingulum_Mid_R | 5 | 20 | 32 |  |  | 5.029 |
| Supp_Motor_Area_R | 8 | 2 | 51 |  |  | 4.899 |
| Precentral_L | -29 | -12 | 65 |  |  | 4.783 |
| Frontal_Sup_L | -24 | 3 | 68 |  |  | 4.677 |
| Cingulum_Ant_L | -3 | 36 | 29 |  |  | 4.639 |
| Frontal_Sup_Medial_R | 5 | 27 | 42 |  |  | 4.405 |
| Frontal_Inf_Tri_R | 43 | 38 | 27 | 0.011 | 1104 | 5.402 |
| Frontal_Mid_R | 27 | 47 | 29 |  |  | 3.993 |
| Insula_R | 59 | -13 | 7 | 0.132* | 525 | 4.520 |
| **Neglectful > Control Mothers (GMV)** |  |  |  |  |  |  |
| Cerebellum_9_R | 21 | -41 | -48 | 0.001 | 4035 | 4.483 |
| Cerebellum_8_R | 17 | -60 | -50 |  |  | 3.830 |
| Cerebellum_Crus2_R | 50 | -69 | -39 |  |  | 3.583 |
| Fusiform_R | 33 | -62 | -18 |  |  | 3.041 |
| **Neglectful < Control Mothers (WMV)** |  |  |  |  |  |  |
| Cluster Frontal_L | -32 | 11 | 29 | 0.000 | 4707 | 5.673 |
|  | -15 | 12 | 48 |  |  | 4.843 |
|  | -8 | 27 | 51 |  |  | 4.555 |
| Cluster Frontal_R | 41 | 21 | 24 | 0.042 | 1805 | 4.503 |
|  | 33 | 29 | 23 |  |  | 4.173 |
|  | 15 | 29 | 32 |  |  | 3.345 |

Note: Whole-brain voxel corrections were applied. All the reported local maxima belong to a significant cluster ( $p$ value FWE corrected < 0.05) with the exception of the probability value signaled with an asterisk indicating that this cluster was significant after a small volume correction (cluster level $p$ value (FWE-corr) = 0.001; peak level $p$ value (FWE-corr) = 0.019). NM: Neglectful mothers; CM: Control mothers. Sup: Superior; Ant: Anterior; Inf: Inferior; Mid: Middle; Supp: Supplementary; Tri: Triangular; R: Right; L: Left.

**Figure 1.**
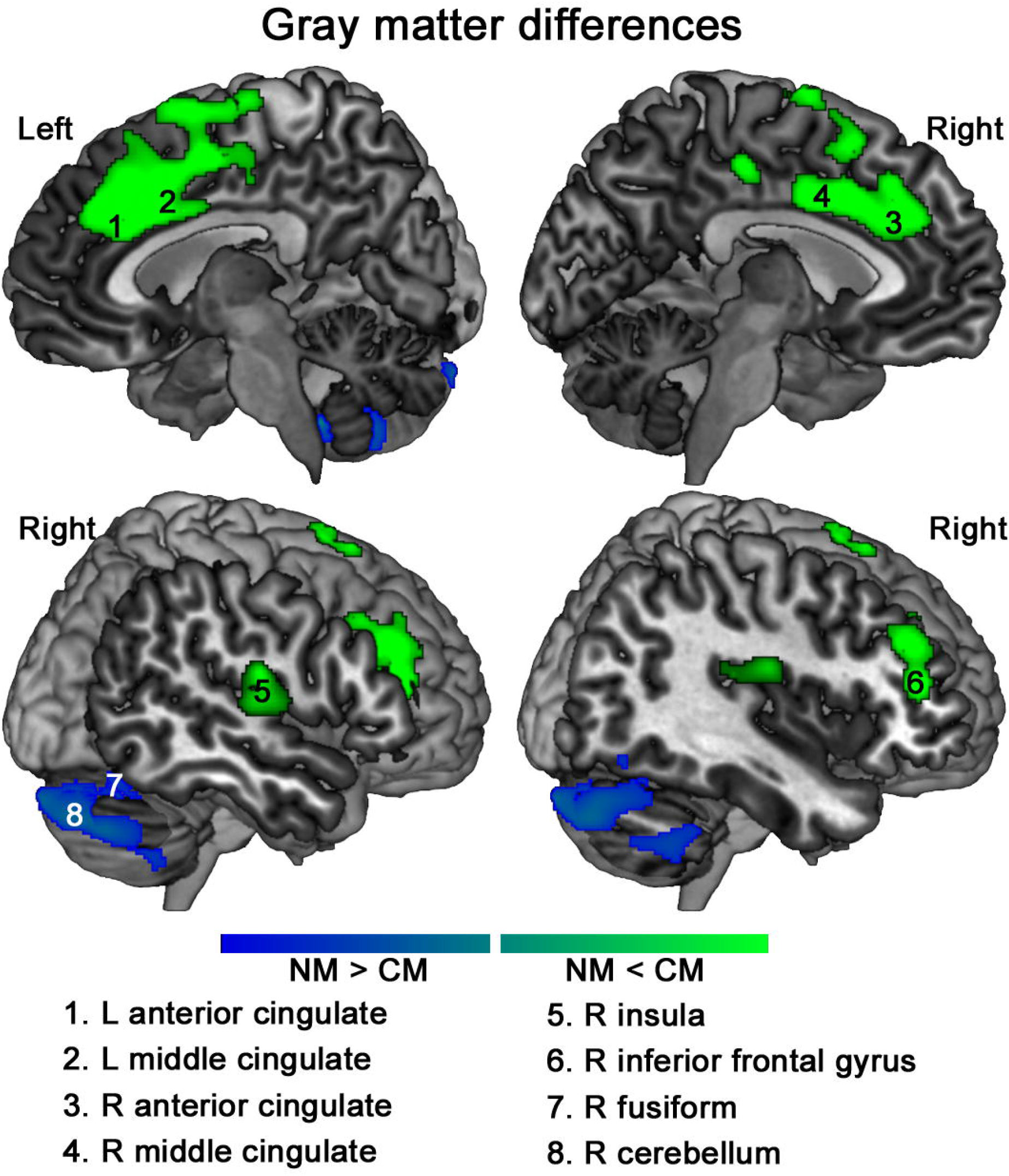
GMV alterations in neglectful mothers in empathy-related regions. GMV reductions in NM as compared to CM were found in the bilateral anterior/middle cingulate cortex, right insula and right inferior frontal gyrus. A reversed pattern of GMV increases in NM was found in the right fusiform and the right cerebellum.

WM analyses showed two frontal bilateral clusters of decreased WMV in NM traversed by the association fiber tracts sensitive to individual differences in empathic concern (See Table 3 and Figure 2). Thus, bilaterally, clusters comprised the inferior fronto-occipital fasciculus (IFOF), the corpus callosum, and the anterior corona radiate (ACR). The left WM cluster also included the anterior thalamic radiation (ATR; Oishi, Faria, Van Zijl, & Mori, 2010). The reduced WMV in the frontal clusters was located adjacent to the two reduced GMV frontal clusters (Figure 3).

**Figure 2.**
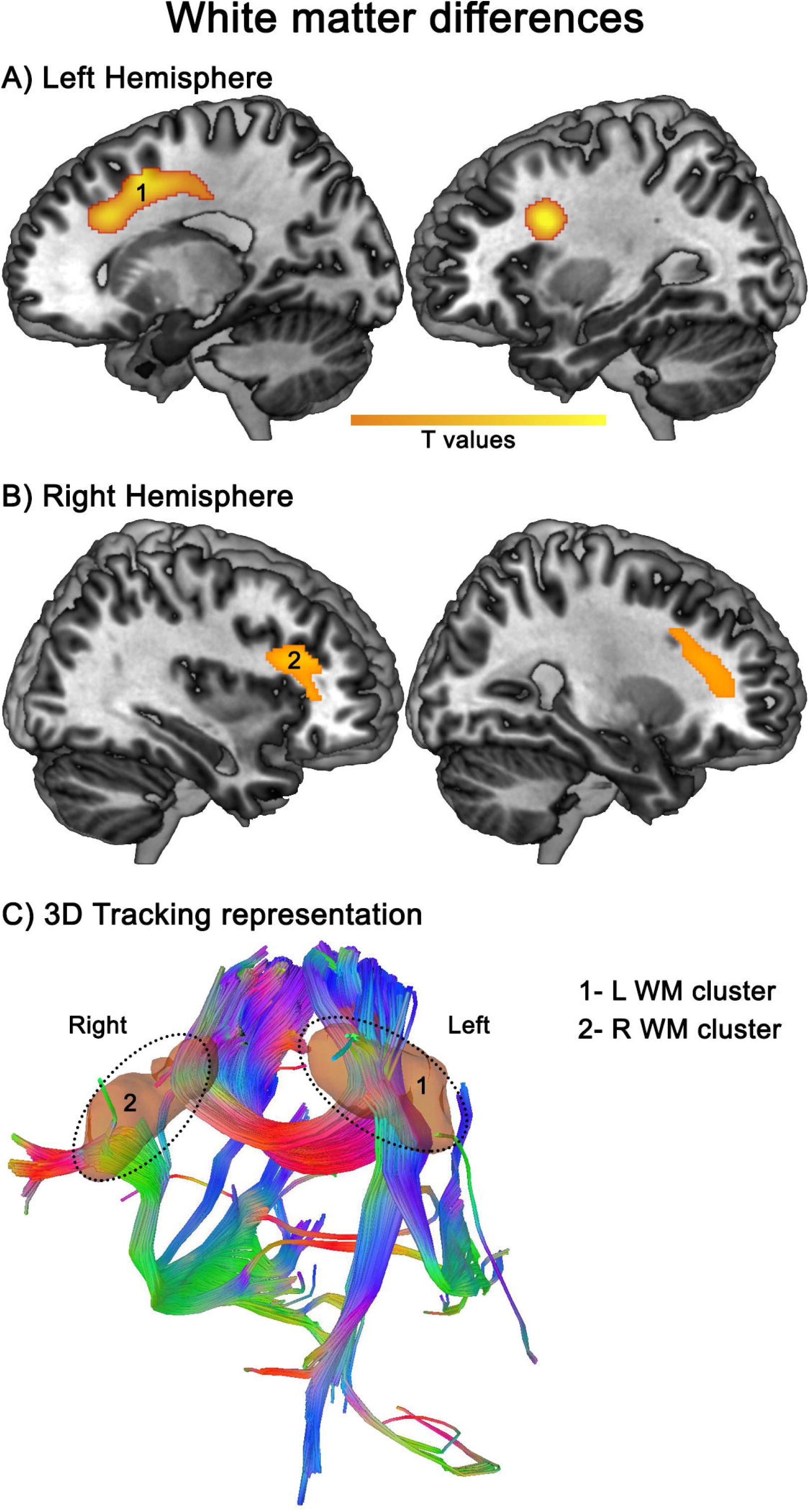
WMV alterations in neglectful mothers in frontal regions. (A and B) WMV reductions in NM as compared to CM were found in two frontal bilateral clusters. (C) 3D tracking representation (see dotted ovals) shows that the right frontal cluster is traversed by the inferior fronto-occipital fasciculus (IFOF), the corpus callosum, and the anterior corona radiate (ACR), whereas the left frontal cluster is traversed by the anterior thalamic radiation (ATR).

**Figure 3.**
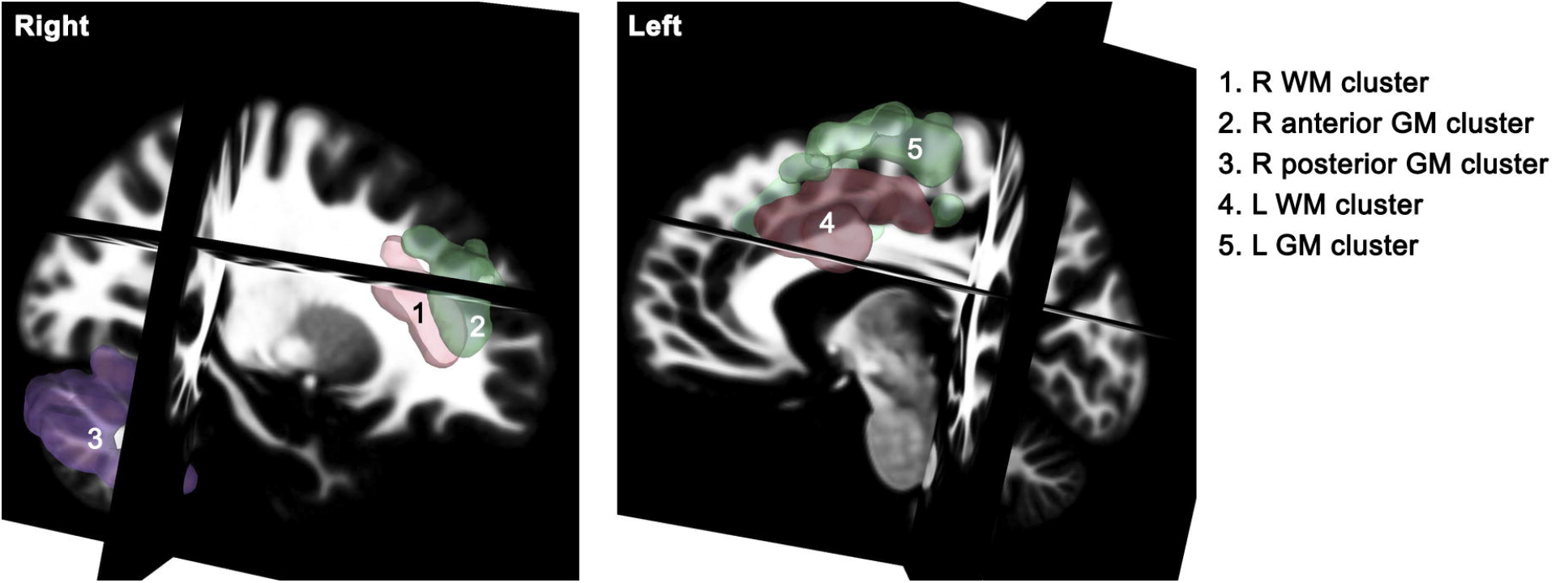
Overlapping of WMV and GMV frontal alterations in neglectful mothers. The reduced WMV in the bilateral frontal clusters were located adjacent to the two reduced GMV in the bilateral frontal clusters.

### Effects of alterations in GMV and WMV on Emotional Availability

To test the hypothesis of whether the altered GMV and WMV predicted mother-child interaction in the play task, we used regression analysis. The five GM areas (ACC/M, R-IFG, Insula, Fusiform_R, and Cerebellum) and the two WM areas (WM_L and WM_R frontal areas) showing altered volume in NM versus CM were used as predictors of the EA. As the majority of NM (17 out of 25) had suffered childhood maltreatment (neglect or physical abuse) as it is usually the case (Mulder, Kuiper, van der Put, Stams, & Assink, 2018), they are more likely to be less sensitive when interacting with their child (Mielke et al., 2016). Therefore, the maltreatment classification (21 maltreated and 27 non-maltreated mothers) was also entered as a variable into the regression to examine its potential contribution to the association of GMV and WMV anomalies with EA. Results showed as significant regressors the GMV of Fusiform_R, the WMV of Frontal_R, and Maltreatment, *F(3,44)* 7.98=, *p* < 0.001), explaining a moderate proportion of the variance on EA (*R*^*2*^ = 0.352; *AdjR*^*2*^ = 0.308). Increased frontal WMV_R predicted higher scores in EA (*β* = 0.26; *t*(44) = 2.08, *p* = .04), whereas increased GMV in Fusiform_R (*β* = −0.24; *t*(44) = −1.94, *p* = .05) as well as belonging to the maltreatment group (*β* = −0.34; *t*(44) = −2.74, *p* = .01) predicted lower scores in EA (Figure 4A & B).

**Figure 4.**
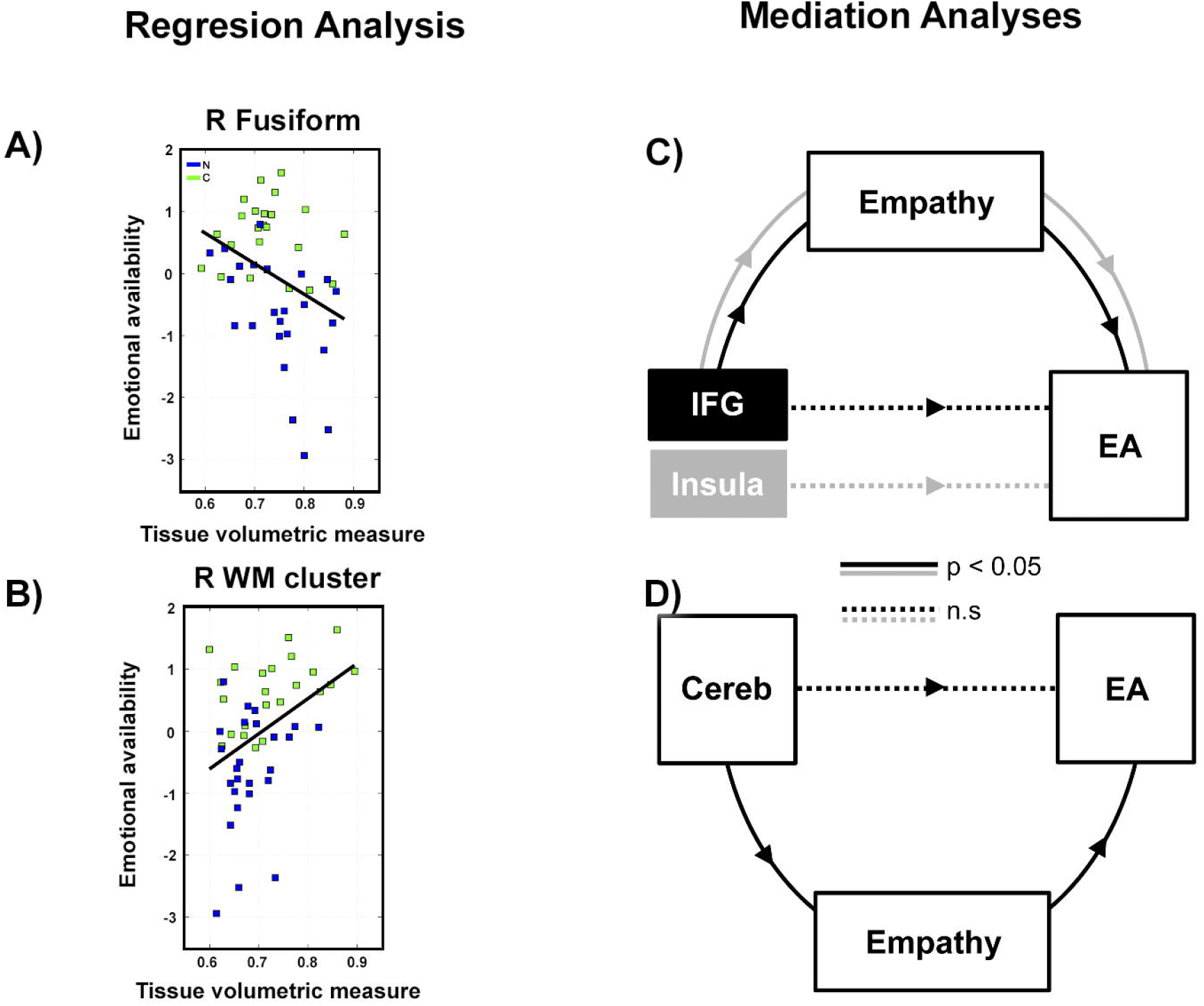
Regression and mediation models of brain differential areas in neglectful mothers on emotional availability (EA). The regression model shows (A) a negative relationship between GMV Fusiform_R and (B) a positive relationship of WMV_R frontal cluster and EA. Mediation models with trait empathy show (C) that Emotional Concern significantly mediated the positive relationship between IFG and Insula volumetric measures and EA, with no direct effects; and (D) that Emotional Concern significantly mediated the negative relationship between Cerebellum volumetric measures and EA, with no direct effects.

### Empathic Concern and Perspective Taking as mediators of the effects of GMV and WMV on Emotional Availability

We performed mediation analyses to further test if those five areas not directly linked to EA in the regression model (ACC/M, IFG_R, Insula_R, Cerebellum, and WM_L) would be related through the mediation of Empathic Concern (EC) and Perspective Taking (PT). For each mediation model with EC, each of the aforementioned areas acted as the independent variable, EC as a mediator, and EA as the dependent variable. Three of the five mediating models showed significant relations. Using a bootstrap resampling procedure, their parameters fell outside the confidence intervals, indicating that the results are not likely to be random. EC mediated in the positive relation of both IFG (Average Causal Mediation Effects, *ACME* = 4.03, *p* = 0.001) and the Insula (*ACME* = 3.39, *p* = 0.03) on EA (Figure 4C), and in the negative relation for Cerebellum (*ACME* = −2.56, *p* = 0.04) on EA (Figure 4D). Neither mediated nor direct significant effects were found for EC in relation to ACC/M and WM_L and EA, and no effect was found for PT for any of the models.

## Discussion

This study revealed alterations in the brain plasticity consisting of a reduced GMV in NM as compared to CM for critical empathy-related areas of the maternal brain including right and left ACC and MCC, right anterior insula, and right IFG, corresponding to those areas showing increased GMV in high empathic adults. The insula-cingulate structures function as an alarm system to infant signals of pain and distress whereas the right IFG is a core region of the mirror neuron system, which enables parents to intuitively resonate with the child’s actions and facial expressions while observing them (Fan, Duncan, de Greck, & Northoff, 2011; Shamay-Tsoory, Aharon-Peretz, & Perry, 2009). NM also showed a pattern of increased GMV in areas crucially involved in emotional face processing, such as the cerebellum (Adamaszek et al., 2017) and right fusiform (Weiner & Zilles, 2016).

The different pattern of GMV changes in NM in the empathy-related areas (decreased frontal areas) and in the viso-emotional processing system (increased occipital areas) is quite compatible with the asymmetric pattern found in EEG rhythms in response to emotional stimuli also found in NM (León et al., 2014): the higher the increases in theta and lower alpha oscillations in response to emotional pictures at occipital sites, the lower the increases of the same bands at frontal sites. A reversed EEG oscillatory pattern was found in CM. Therefore, both GMV and oscillatory patterns in NM would reflect the lower engagement of frontal regulatory processes over the occipital areas in emotional processing. Brain regulatory responding to emotional information is critical for sensitive parenting and for the development of emotion regulation in early child development (Rutherford, Wallace, Laurent, & Mayes, 2015). The current results highlight the relevance of the frontal-occipital distribution of gray matter volume in the context of emotion regulation, as is presumably part of the neuro-cognitive changes associated with sensitive parenting.

In turn, alterations in WMV consisted of a reduction in frontal clusters in NM. The bilateral WM frontal clusters were traversed by fibers of the inferior fronto-occipital fasciculus (IFOF), the corpus callosum, and the anterior corona radiate (ACR), whereas the left WM frontal cluster was traversed by fibers of the anterior thalamic radiation (ATR). These results greatly converge with those found in DTI studies showing reduced integrity in the same tracts in adults with lower scores in Empathic Concern (Parkinson & Wheatley, 2012). Altogether, the GMV and WMV results pointed to an important restriction in volume on a highly interconnected frontal region at the crossroads of viso-emotional pathways, the thalamus with the cortex, and the communication between hemispheres, alterations that may undermine the way mothers “read” the emotional cues to appropriately engage in sensitive interactions with the child.

As expected, the quality of mother-child bonding (EA) was lower in neglectful than in control dyads. Importantly, we obtained evidence of the functional impact of GMV and WMV alterations on the EA, either directly or when introducing differences in Empathic Concern as a mediator. Both increased GMV in right fusiform and decreased WMV in right frontal areas were directly associated with the lower EA, once the negative impact of childhood maltreatment on the mothers had been teased out (Mielke et al., 2016). The frontal anomalies found in WMV seem to correspond to the lower structural connectivity in the IFOF obtained in a DTI study in NM, and also related to poor mother-child bonding (Rodrigo et al., 2016). Although NM scored lower on both Empathic Concern and Perspective Taking than CM (León et al., 2014; Rodrigo et al., 2011), increased volume in emotional empathy-related areas such as insula and IFG predicted higher EA only mediated by the Empathic Concern. In turn, increased cerebellum volume mediated by Empathic Concern predicted a lower EA, suggesting its implication in emotional empathic processes (Adamaszek et al., 2017). These brain-behavior associations could be used in the future as a biological indicator of dysfunctional mother-child interactive behavior.

The GM areas affected in NM associated with EA as well as the unique mediating role of Empathic Concern in this relation suggest that emotional empathy-related areas, responsible for the automatic perception of distress signals and involved in embodied simulation and sympathy with others’ emotions, played a distinctive, prominent role in the caregiving network. NM’s alterations in the emotional empathy-related areas may disturb the so-called “intuitive parenting” (Papoušek & Papoušek, 1987) that provides a fast response to the child that is crucial to child attachment (Altenhofen et al., 2013). A convergent finding is that mothers’ caregiving behavior is particularly driven by the emotion-processing network while fathers when acting as primary caregivers, exhibit activity in both the emotional and the cognitive empathy areas (Abraham et al., 2014). In turn, the volumetric reduction in NM in the anterior/middle cingulate cortex presumably related to cognitive empathy (Eres et al., 2015) was not associated with EA, confirming a less prominent role of the cognitive empathy-related areas in neglectful caregiving.

Our present results converged with those found with EEG and DTI (León et al., 2014; Rodrigo et al., 2016, 2011) in defining some of the alterations underlying maternal neglect. A paradoxical result is the increased GMV in cerebellum and fusiform areas, where an activation reduction was found in these areas in response to crying faces in an fMRI study with the same mothers (León et al., in press). However, a greater convergence (reductions both in GMV and activation to infant crying faces) was found in frontal and cingulate areas. More research is needed to envision how the neural organization is altered in these mothers at structural, functional, and connectivity levels and to elucidate the potential connections between levels. In the same line, larger samples with an orthogonal design crossing NM and CM with the pathological conditions as well as other risk factors (e.g., their own childhood maltreatment or epigenetic changes) will allow us to determine —and not only control for — their contribution to the neural alterations associated with maternal neglect. Finally, future studies should use a longitudinal design to fully ascertain during the postpartum period a lack of or abnormal changes in the maternal plasticity in NM and their causal relationships to EA.

## Conclusion

In conclusion, the “neglectful” brain seems not to have benefited from the typical maternal adaptations to sensitive caregiving. Specifically, the increased GM volume in emotional empathy areas, not found in NM, seems to play a prominent role in maternal brain plasticity well after the postpartum period. GM alterations in NM may affect their “intuitive parenting” capacities that enable more automatic, fast and emotional responding to the needed child. The mothers’ poor emotional attunement to the others’ signal of distress may be imitated by the neglected child, leading to the poor interactive bonding exhibited by our neglectful dyads. In turn, the establishment of positive mother-child interactions also requires the increased WM volume in a highly interconnected frontal region, not found in NM, that seems to exert regulatory control over the processing of viso-emotional information. This is a new proposal that deserves closer examination by means of connectivity analyses. Altogether, neglectful mothering is characterized by GM and WM volumetric alterations, affecting both the automatic and elaborative processes that may be engaged when establishing mutual emotional bonds with the child. Prevention and intervention strategies training mothers to manage their own emotions when faced with their distressed infant and to enhance their emotional empathic responding are necessary to ensure the neural equipment that maximizes infant survival and wellbeing.

## Acknowledgments

This work was supported by the Spanish Ministry of Economy and Competitiveness and the European Regional Development Fund under Grant PSI2015-69971-R to M.J.R. & I.L. We thank Dr. Alejandro Pérez for his comments on the manuscript. We also thank the Health and Social services staff and all the mothers and their infants who participated in this study.

## APPENDIX

1) Social workers reported on a series of risk indicators (presence: 1; absence: 0) that are commonly used to assess maternal neglect. *History of abuse/neglect* refers to whether mothers have suffered childhood maltreatment (either abuse or neglect) in their own history (scoring 1); *Intimate partner conflict* refers to whether mothers are experiencing overt conflictive relationships with their partner (scoring 1); *Chronic physical illness* refers to whether they are currently experiencing poor health conditions permanently or very frequently (scoring 1); *Poor household management* refers to whether the home is dirty and/or untidy, with irregular meals and/or dirty clothing (two is enough for scoring 1); *Disregard health/education needs* refers to lack of or discontinuous medical checks, irregular vaccines, and/or poor support for learning (two is enough for scoring 1); *Disregard emotional/cognitive needs* refers to poor attention to the child’s emotional expressions and/or lack of response to infant curiosity (one is enough for scoring 1); *Rigid/inconsistent parental norms* refers to an application of rules without taking into account the childrearing situations and/or arbitrary changes to norms applied to the same situations (one is enough for scoring 1).

**Table A1.** Collinearity indexes between the Group (as a dichotomic variable) and the psychopathological conditions, both as a factor PD and separately as individual disorders (within brackets are the cutoff values for non-collinearity).

|  | VIF<br>(<10) | TOL<br>(>0.30) | CN<br>(<10) | Shared<br>Variance<br>(<0.50) |
| --- | --- | --- | --- | --- |
| **Factor score**<br>**"Psychiatric**<br>**Disorder"** | **1.80** | **0.56** | **2.23** | **0.44** |
| Major<br>Depressive<br>Disorder | 1.23 | 0.81 | 1.58 | 0.18 |
| Hypo/Manic<br>Episode | 1.34 | 0.75 | 1.70 | 0.24 |
| General Panic<br>Disorder | 1.54 | 0.64 | 1.92 | 0.33 |
| Generalized<br>Anxiety<br>Disorder | 1.30 | 0.77 | 1.69 | 0.23 |
| Antisocial<br>Personality | 1.42 | 0.70 | 1.74 | 0.25 |

Note: VIF: Variance Inflation Factor, TOL: Tolerance, CN: Condition Number.

We assessed the potential collinearity (CL) calculating three well-known indexes of CL: the Variance Inflation Factor (VIF), the Tolerance (TOL), and the Condition Number (CN). We calculated them for the overall factor PD with the Group, as well as separately for each psychiatric disorder that survived the Bonferroni test in the group comparisons (Table A3). We also measured the shared variance of the psychiatric variables with Group (last column). According to the literature, the general criterion for non-collinearity is a Variance Inflation Factor (VIF) less than 10, a Tolerance greater than 0.30, a Condition Number less than 10, and a shared variance less than 0.50 (Belsley, 1991; Belsley et al., 2005; Kovács et al., 2005; Tabachnick & Fidell, 2007). Our results (see Table A3) showed that all the values fall below the corresponding cutoffs for collinearity. That made it possible to include the PD as a covariate together with Group in the SPM model, to control as much as possible its effect on the activation results.

**Table A2.** Inter-rater reliabilities and one-factor component loadings of the Emotional Availability Scales.

|  | Kappa<br>coefficients | Component<br>loadings |
| --- | --- | --- |
| Sensitivity | 0.94 | 0.871 |
| Structuring | 0.90 | 0.867 |
| Nonintrusiveness | 0.87 | 0.857 |
| Nonhostility | 0.92 | 0.762 |
| Responsiveness | 0.92 | 0.799 |
| Involvement | 0.86 | 0.783 |

For the emotional availability score, the mother-child interaction was videotaped at home, in the context of mother-child free play, at the moment when the family received a toy as a gift for participation in the study. Mothers were instructed to use the toy and play with the child as they usually do. Ratings from the videos were based on the Emotional Availability Scale, which operationalizes four aspects of parental behavior: *Sensitivity* (9 points) - the parent shows contingent responsiveness to child signals; *Structuring* (5 points) - the parent appropriately facilitates the child’s play; *Non-intrusiveness* (5 points) - the parent is able to support the child’s play without being overdirective and/or interfering; *Non-hostility* (5 points) - the parent is able to behave with the child in a way that is not rejecting or antagonistic. The scale also measures two aspects of child behavior: *Responsiveness* (7 points) - the child’s ability and interest in exploring on his or her own and in responding to the parent’s bids; *Involvement* (9 points) - the child’s ability and willingness to engage the parent in interaction (Table A4). To obtain a more simple structure of the six standardized scales, a Principal Component Analysis was performed. The result yielded a single factor structure: KMO = 0.82, Eigenvalue = 4.34, with an explained variance of 72%.

